# Mechanistic insights from replica exchange molecular dynamics simulations into mutation induced disordered-to-ordered transition in Hahellin, a βγ-crystallin

**DOI:** 10.1101/475277

**Authors:** Sunita Patel, Bal Krishnan, Ramakrishna V. Hosur, Kandala V. R. Chary

## Abstract

Intrinsically disordered proteins (IDPs) form a special category because they lack a unique well-folded 3D structure under physiological conditions. They play crucial role in cell signaling, regulatory functions and responsible for several diseases. Although, they are abundant in nature, only a small fraction of it has been characterized till date. Such proteins adopt a range of conformations and can undergo transformation from disordered-to-ordered state or vice-versa upon binding to ligand. Insights of such conformational transition is perplexing in several cases. In the present study, we characterized disordered as well as ordered states and the factors contributing the transitions through a mutational study by employing replica exchange molecular dynamics simulation on a βγ-crystallin. Most of the proteins within this superfamily are inherently ordered. However, Hahellin, although a member of βγ-crystallin, it is intrinsically disordered in its apo-form which takes a well-ordered βγ-crystallin fold upon binding to Ca^2+^. It is intriguing that the mutation at the 5^th^ position of the canonical motif to Arg increases the domain stability in several ordered microbial βγ-crystallins with concomitant loss in Ca^2+^ binding affinity. We carried out similar Ser to Arg mutations at 5^th^ position of the canonical motif for the first time in an intrinsically disordered protein to understand the mechanistic insights of conformational transition. Our study revealed that newly formed ionic and hydrogen bonding interactions at the canonical Ca^2+^ binding sites play crucial role in transforming the disordered conformation into ordered βγ-crystallin.

**Author summary:** Intrinsically disordered proteins lack a unique ordered 3D structure under physiological condition. Although, they are abundant in nature, only a small fraction of these proteins has been characterized till date due to adaptation of multiple conformations and methodological limitation. βγ-crystallins are inherently ordered, however recently a small number of proteins within this superfamily have been identified as intrinsically disordered protein. Hahellin is one such protein which is intrinsically disordered in its apo-form but takes a well-ordered βγ-crystallin fold upon binding to Ca^2+^. In the present study, we decipher the underlying mechanism of disordered-to-ordered transition in Hahellin by mutations, employing replica exchange molecular dynamics simulations. Earlier experimental studies reported an increase in stabilization of the ordered βγ-crystallion upon mutation to Arg at 5^th^ position of the canonical Ca^2+^ binding motifs, N/D-N/D-X_1_-X_2_-S/T-S. We performed similar Ser to Arg mutation in an intrinsically disordered Hahellin to get the mechanistic insights of the conformational transition in the absence of Ca^2+^. Our study revealed that several newly formed ionic and hydrogen bonding interactions contributed by the mutant residues are responsible for both intra- and inter-motif rigidification, resulting in overall stability of βγ-crystallin domain.

## Introduction

Proteins lacking a stable 3D structure under physiological conditions are classified as intrinsically disordered proteins (IDPs). Intrinsically disordered proteins although are prevalent in nature only a small fraction of these proteins have been characterized till date [1, 2]. This is because of several challenges associated with them, which could be due to multiple conformations attainable by the same polypeptide chain and/or methodological limitation. Until, recently, IDPs, found in all kingdoms of life [1] were unexplored, and their existence and significance have now been realized by innumerable examples [3, 4]. Recently, the IDPs have been shown to be crucial in many signal transduction pathways and regulatory functions [2, 5]. They have also been shown to be responsible for several human diseases such as Alzheimer, Parkinson, Huntington, type II diabetes mellitus, cancer, cardiovascular disease, cataract and others [3, 6-9]. They are also recently surmised to be an attractive target for novel therapeutics [10]. Thus, considering the involvement of IDPs in a diverse array of biological functions, understanding their structural and dynamical properties is of great significance.

IDPs are highly flexible and can exist in several interconverting states under physiological conditions [4, 11, 12]. They often undergo large conformational changes through post-translational modifications or upon binding to a ligand. The ligand can be as small as a metal ion or a biomolecule such as membrane, DNA or protein. The conformational change can be either from disordered-to-ordered state or vice-versa [12-14]. The disordered-to-ordered transition, most often is a coupled event of folding and binding [12-14]. Understanding the molecular basis of disordered-to-ordered transition is therefore vital to decipher the responsiveness of these agents to modulate the cellular functions.

βγ-crystallins are inherently ordered proteins and they are well known for their intrinsic domain stability as observed in the eye-lens crystallins which provides transparency to the eye-lens and present throughout the lifespan of an organism. βγ-crystallins includes proteins from diverse origin such as archaea, bacteria and eukaryotes. [15]. Most of the archaeal and bacterial crystallins such as Protein S, M-crystallin, Clostrillin are ordered well-folded and undergo further stabilization upon binding to Ca^2+^ while vertebrate crystallins are highly stable, but display low affinity for Ca^2+^ [16, 17]. Such structural inconsistency arose curiosity. Although, βγ-crystallins includes proteins which are highly ordered, recently a number of proteins in this superfamily have been identified as intrinsically disordered. For example, Hahellin, along with a couple of other intrinsically disordered βγ-crystallins such as Caulollin [18] and Yersinia crystallin [19], represent extreme Ca^2+^ dependency for their structural stability [20]. Thus, many proteins in this superfamily bind to Ca^2+^ and undergo further stabilization while others do not bind to Ca^2+^, but still get similar stabilization. The significance of such states is yet to be understood.

Hahellin is a protein domain encoded by the genome of a marine bacterium *Hahella chejuensis* which grows under extreme conditions. It has sequence signature of a βγ-crystallin. Hahellin, upon binding to Ca^2+^, undergoes a drastic conformational transition and acquires a βγ-crystallin fold [20]. A typical βγ-crystallin fold consists of two consecutive Greek keys motifs with double-clamp architecture having two canonical N/D-N/D-X_1_-X_2_-S/T-S, Ca^2+^ binding sites [20-24]. The well-folded 3D structure of Ca^2+^-bound Hahellin (holo-Hahellin) and its conformational heterogeneity and dynamics were studied by NMR spectroscopy [20, 25]. Subsequently, in another study, a combined use of NMR and REMD simulation was used to characterize the intrinsically disordered states of Hahellin (apo-Hahellin) which revealed heterogeneous mixture of near-native and far-native conformations [26]. Such a conformational ensemble was attributed to the repulsive interactions of the negatively charged residues located in the Ca^2+^ binding sites in the absence of metal ion. Besides, a few prior studies on ordered βγ-crystallins delineate that the residue at the 5^th^ position of the canonical Ca^2+^ binding sites (N/D-N/D-X_1_-X_2_-S/T-S) which is normally occupied by a Ser or Thr when mutated to Arg, increases the domain stability [17, 22]. However, the underlying mechanism of gain in stabilization upon mutation is still baffling.

Therefore, to understand the effect of mutation on the intrinsically disordered Hahellin structure and dynamics, we mutated the two key Ca^2+^-ligating Ser residues into Arg at each canonical motif, individually and together, and examined the consequences. Although, such mutations have been studied on few ordered βγ-crystallins, so far, no study has been carried out on an intrinsically disordered protein which belongs to the βγ-crystallin superfamily. In the present study, we observed that the mutation at the 5^th^ position of both the canonical motifs (N/D-N/D-X_1_-X_2_-S/T-S) of Hahellin from Ser to Arg reduces the conformational heterogeneity and promotes homogeneity into an ordered βγ-crystallin without the necessity of Ca^2+^, which otherwise in the absence of Ca^2+^ imparts intrinsically disordered structure. The newly formed ionic and hydrogen bonding interactions at the canonical Ca^2+^ binding sites causes intra- and inter-motifs rigidification and thus play very crucial role in transforming the disordered conformation into a ordered βγ-crystallin.

## Results

### Suppression of conformational heterogeneity in Hahellin mutants

The REMD simulation on the Hahellin-wt shows widely distributed C^α^ RMSDs ranging from 2 to 12 Å [26] (Figure 1). The C^α^ RMSD is determined with respect to the well-folded holo-Hahellin. Figure 1 shows eight peaks which is indicative of heterogeneous populations. Some of these peaks are overlapping, suggesting co-existence of more than one conformational types. The conformations having smaller values of C^α^ RMSD are similar to holo-Hahellin and therefore represent near-native conformations, while those having higher values of C^α^ RMSD represent far-native conformations, as reported earlier in our study [26]. In the present study, the REMD simulations on the three mutants (Hahellin-S41R, Hahellin-S80R and Hahellin-S41R-S80R) show three distinct distributions of C^α^ RMSD (see Figure 1). The C^α^ RMSD of Hahellin-S80R ranges from 2 to 8.2 Å and has three distinct populations centred around 3.2, 5.8 and 8 Å. Two out of the three distributions are largely populated suggesting existence of two major conformational forms. In the second mutant, Hahellin-S41R, the C^α^ RMSD spread is relatively smaller and it ranges from 2 to 6 Å. It shows three overlapping populations centred around 3.2, 4.2 and 5.8 Å suggesting, a major fraction of the conformations are closer to holo-Hahellin compared to that observed in Hahellin-S80R. On the other hand, the third mutant (double mutant) Hahellin-S41R-S80R shows further reduction in the C^α^ RMSD distribution as compared to the rest three simulations. The C^α^ RMSD shows a single distribution having much smaller spread ranging from 2 to 4.2 Å with a peak which is centred around 3 Å. Taken together, the C^α^ RMSD analysis reveals that the Hahellin-wt exhibits diverse conformational forms and the heterogeneity is reduced in the single mutants (Hahellin-S80R and Hahellin-S41R) and is further reduced substantially in the double mutant (Hahellin-S80R-S41R) leading to single homogeneous conformational form.

**Figure 1.**
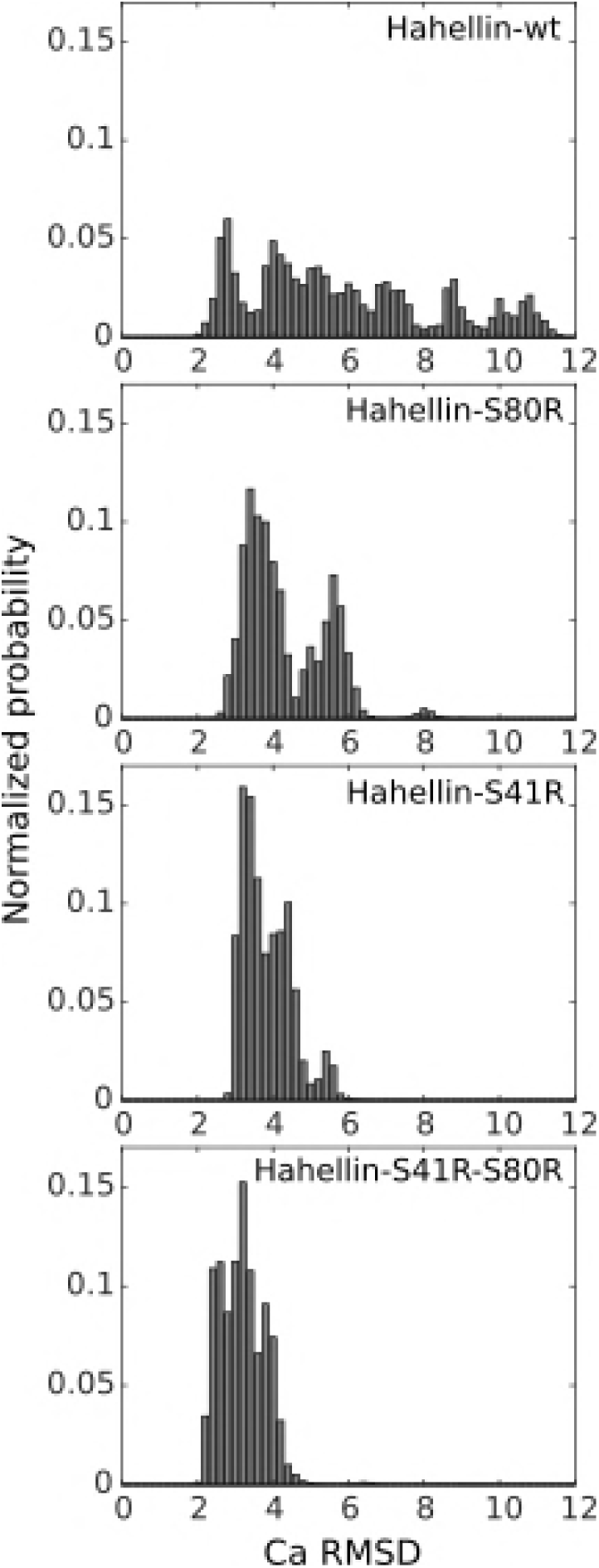
Histogram distributions of C^α^ RMSD w.r.t. holo-Hahellin were determined for Hahellin-wt, Hahellin-S41R, Hahellin-S80R and Hahellin-S41R-S80R simulations.

### Reduction in conformational subspaces of Hahellin mutants on the free energy landscape

The backbone Rg and C^α^ RMSD values are used as conformational coordinates to construct the free energy landscapes, as shown in Figure 2. Hahellin-wt displays large conformational subspace with ten free energy minima, suggesting the ensemble represents a heterogeneous population. Out of the ten minima, three (M1, M2 and M3) are highly populated with Rg values ranging from 12 to 13 Å and RMSD values ranging from 2 to 6 Å. The topologies corresponding to these minima are similar to holo-Hahellin and represent near-native conformations. On the other hand, the remaining minima are relatively less populated (M4 to M10) and are higher in free energies, depicting far-native conformations. In the case of Hahellin-S80R, the constructed Rg-RMSD conformational subspace exhibited five free energy minima, out of which two minima (M1 and M4) have holo-hahellin like β-sheet topology and thus correspond to near-native conformation while the other three minima (M2, M3 and M5) represent far-native conformations (Figure 2B). The minima M3 and M5 with far-native conformations show relatively larger values of Rg, 12.5 and 13.2 Å and higher RMSD, 6 Å and 8 Å respectively while M2 has smaller Rg (12 Å) like M1 but has different RMSD (5 Å). On other hand, for Hahellin-S41R shows four minima in the Rg-RMSD landscape and all of which are compact with Rg values less than 12.5 Å having holo-form like topology except the loop regions which adopt different conformations in these minima. On other hand in the double mutant, Hahellin-S41R-S80R, we noticed reduction in the Rg-RMSD conformational subspace compared to Hahellin-wt and Hahellin-S80R but has similar subspace like that of Hahellin-S41R (Panels C and D Figure 2). However, instead of four minima, Hahellin-S41R-S80R shows only one minimum (M1) which suggests sampling of homogeneous near-native conformations. It has Rg value of 12 Å similar to Ca^2+^ bound Hahellin derived from NMR (pdb id 2KP5) and 3 Å Cα;RMSD with respect to the same NMR-derived holo-Hahellin. Taken together, there is substantial reduction in Rg-RMSD conformational subspace with single minimum for Hahellin-S41R-S80R simulation, hinting that Arg mutations at both canonical sites in place of Ser restricted the conformational sampling into a single minimum.

**Figure 2.**
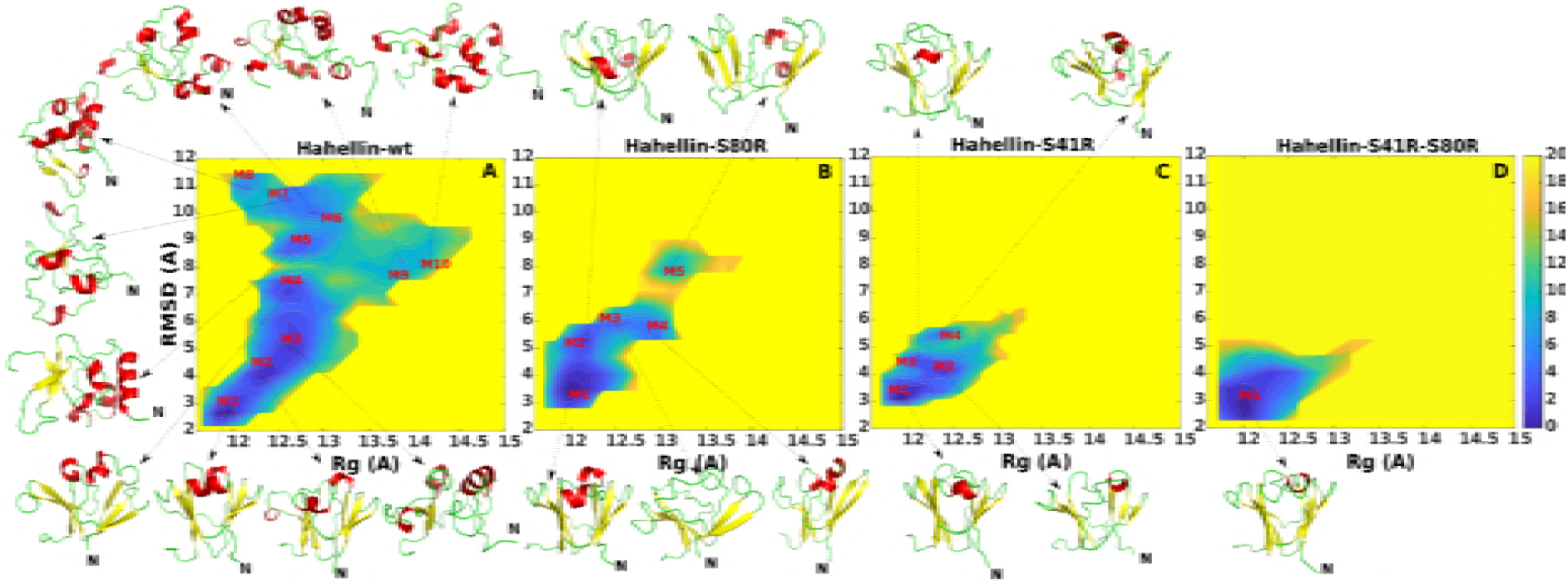
The free energy landscape constructed using backbone Rg and C^α^ RMSD as conformational parameters. The free energy spread in each simulation indicates the corresponding conformational sampling. The sampled regions in the free energy landscape are shown in contours and the free energy values are indicated by colour bar. The conformation corresponding to each free energy minimum are shown by arrow.

### Network layouts show disordered-to-ordered transition upon mutation

Network layouts are constructed for each simulation keeping identical pairwise RMSD cutoff as described in Methods (Figure 3). The Hahellin-wt network layout resulted in ten clusters, while the Hahellin-S80R gave rise to only three clusters and both Hahellin-S41R and Hahellin-S41R-S80R showed further reduction in the number of cluster and resulted into a single cluster (Figure 3). So, there is a progressive decrease in the number of clusters from the wild type protein to single mutants and then in double mutant. In Hahellin-wt, the highly populated cluster has 61.6% of population and its topology is similar to holo-Hahellin while the remaining clusters are less populated and have disordered structure (Figure 3). The percentage of native contact in the Cluster 1 centroid structure (C1) is 78.3%, while for the rest of the clusters, the native contact percentages are less than 36% indicating that Hahellin-wt has heterogeneous mixture of both near-native and far-native conformations, as reported earlier in our study (Figure 5A)[26]. The disordered structures show an increase in helical and loop propensity while retaining some β-strand propensity (Figure S1). The β-strand propensity is observed mostly for the β-strands corresponding to the second Greek key (residue 46 to 90), while the first Greek key (residue 1 to 45) is found to be highly disordered (Figure 4). The Hahellin-S80R network layout shows one dominant cluster (C1) with 96.6% of population and two small clusters (C2 and C3) having 2.8% and 0.6% of population (Figure 3). The dominant cluster (C1) exhibited topology similar like that of holo-Hahellin. Its centroid structure C1, retains ~70.9% of the native contacts while the other two (C2 and C3) have 44% and 32.8% of the native contacts (Figure 4A). For both Hahellin-S41R and Hahellin-S41R-S80R REMD simulations, the resultant network layouts show a single cluster with 100% of population each. They possess 65.7% and 75.4% of the native contacts, respectively, which indicate holo-Hahellin like fold. The centroid structures in both simulations retain similar double Greek key β-sheet topology like that of Ca^2+^ bound Hahellin (Figure 4A). So, a transition from heterogenous mixture to homogenous form is brought about upon mutation to Arg at the 5^th^ position of the canonical Ca^2+^ binding motifs.

**Figure 3.**
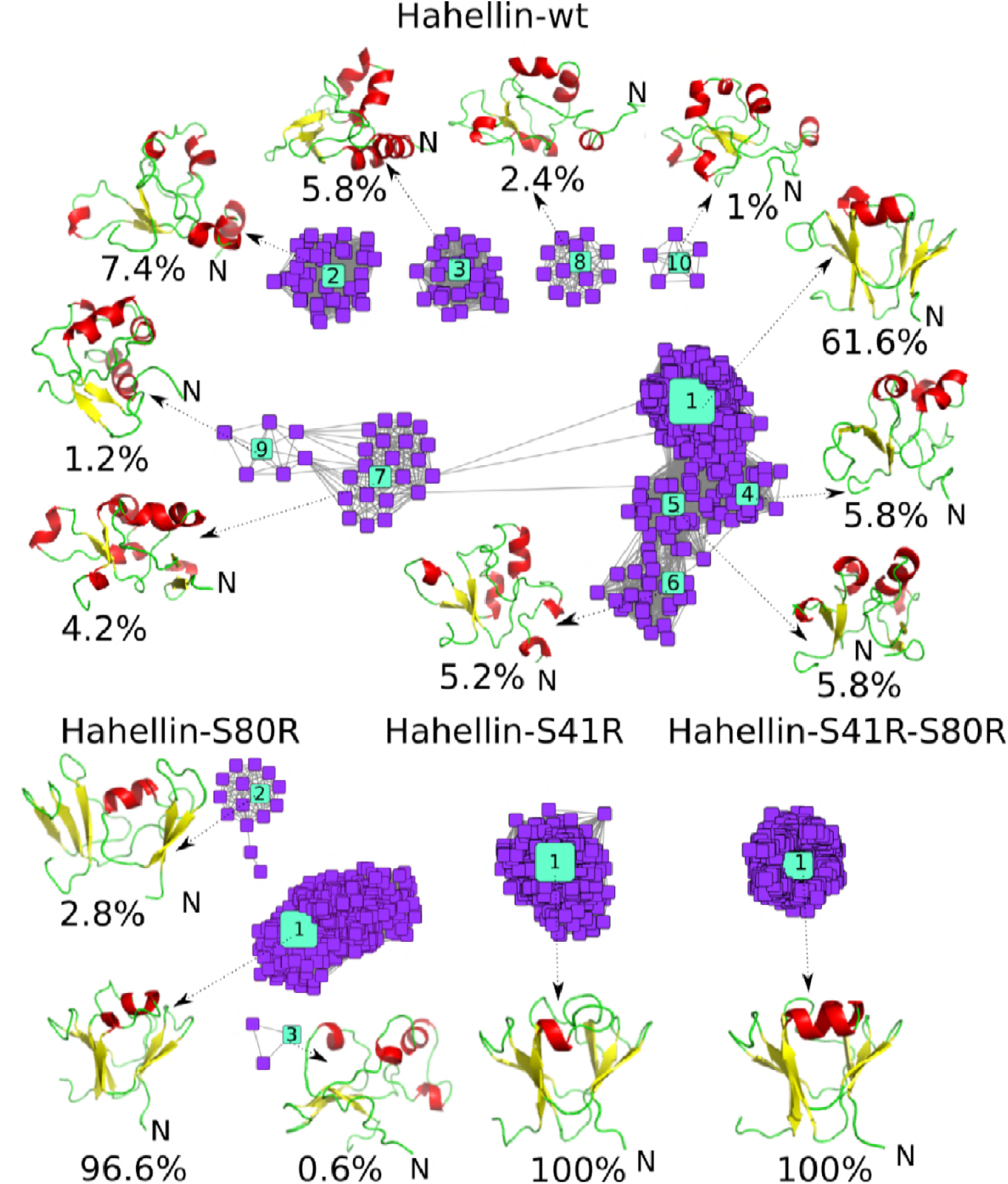
Network layouts of Hahellin-wt, Hahellin-S41R, Hahellin-S80R and Hahellin-S41R-S80R REMD simulations. An identical pairwise RMSD cut-off of 5 Å is used to generate all the layouts. The network layouts are made considering 500 structures obtained at regular interval since this is a memory demanding analysis. The centroid structure corresponding to each cluster are shown by dotted arrow. The cluster percentages are indicated below each centroid structure.

**Figure 4.**
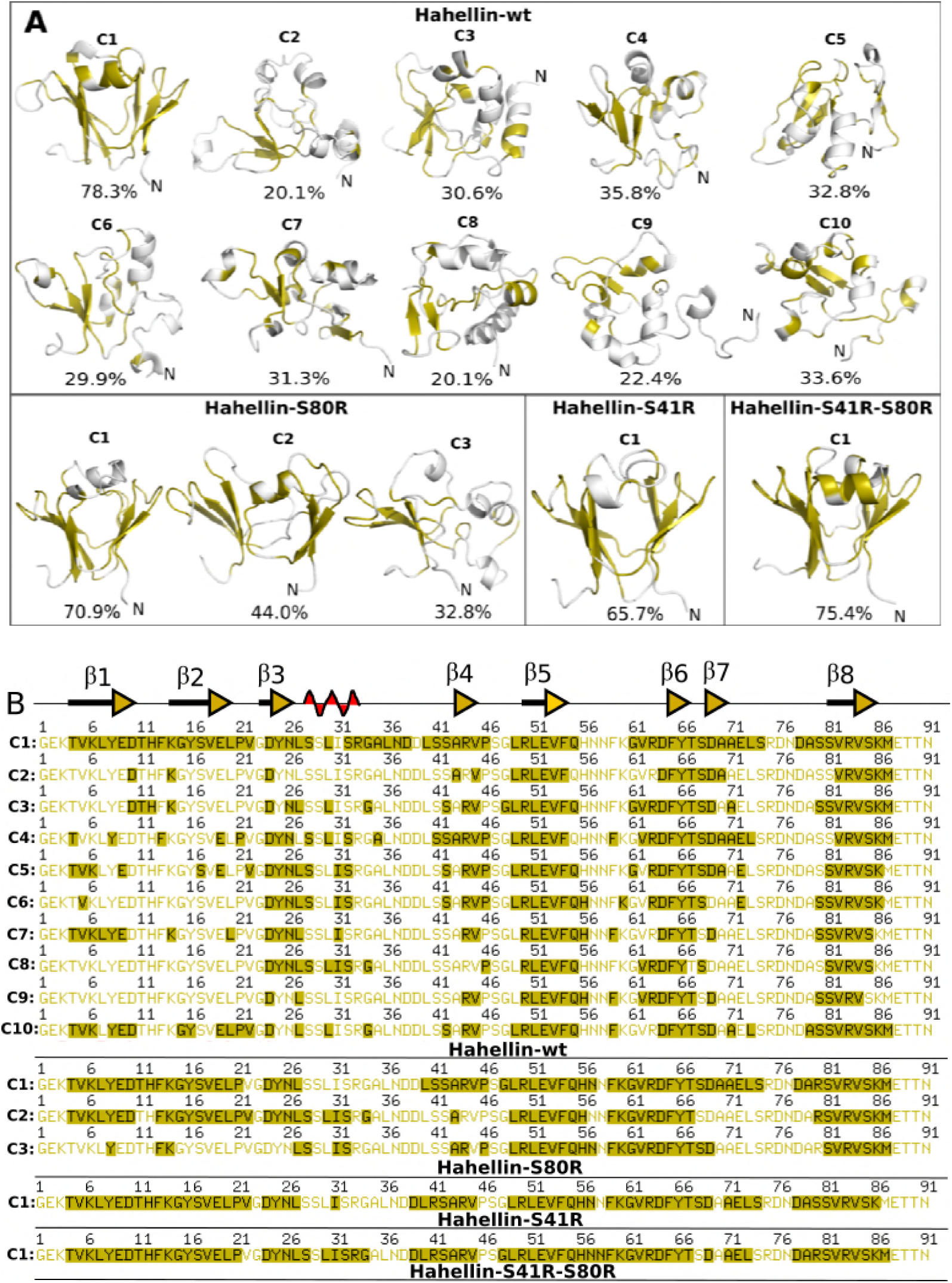
(A) Percentage of native contacts shown for the centroid structure of the cluster with respect to holo-Hahellin. Residues retaining native contacts are mapped on the centroid structure with green colour. (B) The same residues are shaded in green in the amino acid sequences which show the secondary structural information of holo-Hahellin on the top of the sequence.

**Figure 5.**
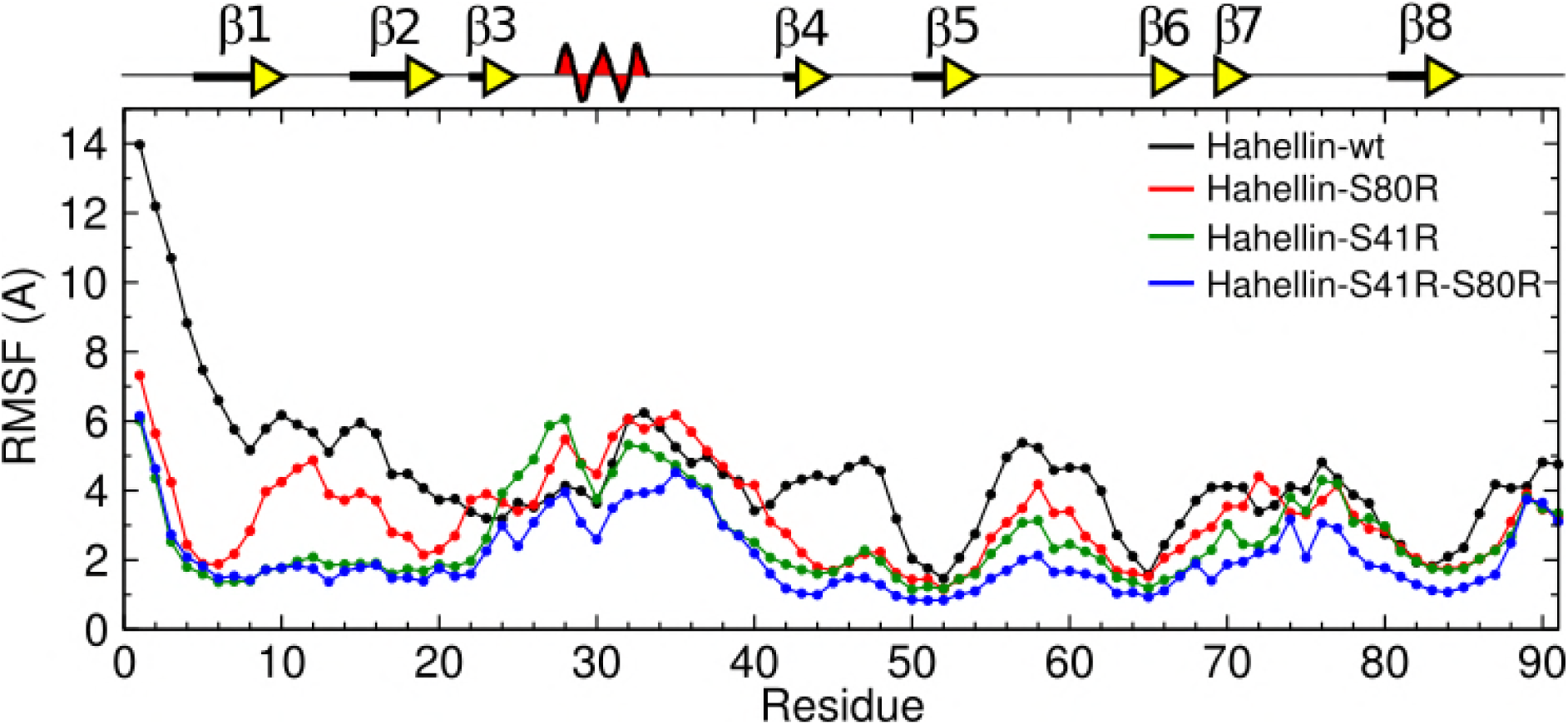
Root mean square fluctuation of C^α^ atoms of each residue in Hahellin-wt, Hahellin-S41R, Hahellin-S80R and Hahellin-S41R-S80R REMD simulations. Secondary structures of NMR derived holo-Hahellin are indicated in the top panel.

### Root-mean-square-fluctuations show differential flexibility

Figure 5 shows the residue-wise root-mean-square-fluctuations (RMSFs) along the polypeptide chain for all the simulations. The terminal ends are highly flexible in all simulations with higher flexibility observed for N-terminal end. Among all, Hahellin-wt displays highest flexibility followed by Hahellin-S80R and then Hahellin-S41R and Hahellin-S41R-S80R (Figure 5). The flexibility in Hahellin-wt is throughout the polypeptide chain except for the β5, β6 and β8-strands located in the second Greek key. Similar to Hahellin-wt, the flexibility in Hahellin-S80R is more for the first Greek key compared to the second Greek key. The regions corresponding to β1-strand in the N-terminal Greek key and β4, β5, β6 and β8-strands in the C-terminal Greek key of Hahellin-S80R show lower values of ~2 Å RMSF, indicating least flexibility for these regions (Figure 5). Compared to Hahellin-wt in Hahellin-S80R, β1 and β4-strands are more rigid. In Hahellin-S41R, the RMSF value further reduced to ~1.8 Å for most of the regions except for the inter-βstrand loops (β5 and β6), (β7 and β8) and short helical regions. The double mutant Hahellin-S41R-S80R shows lowest RMSF (~1.6 Å) all over the polypeptide chain except for the short helix and the inter-βstrand loop region between β7 and β8-strands. It is noteworthy to mention here that the first Greek key is inherently more disordered than the second Greek key in Hahellin-wt. Therefore, mutation at the N-terminal motif exerts overall stabilizing effect. The RMSF profile thus alludes the presence of some interactions in the first Ca^2+^ binding motif of the mutants, Hahellin-S41R and Hahellin-S41R-S80R, which are absent in Hahellin-S80R and Hahellin-wt proteins. These interactions could provide the required stability for the structural integrity of the βγ-crystallin fold.

### Decrease in negative surface charge potential upon mutation

As shown in the Figure 6, we determined the surface charge potential for the centroid structure of the most populated cluster corresponding to each simulation. The NMR derived holo-Hahellin structure is compact and shows a dense red patch in the Ca^2+^ binding loop regions indicating cluttering of negative charge potentials on the surface. These negative charge potential regions favors interaction with positively charged Ca^2+^ ions and contribute to the overall stability of βγ-crystallin fold in the holo-form. The Hahellin-wt on the other hand, is devoid of Ca^2+^ and the centroid structure of the most populated cluster (taken from Figure 3) is loose and less compact than NMR-derived Hahellin. It shows broad distribution of negative surface charge potential on the Ca^2+^ binding loop regions as shown in the Figure 6, suggesting highly destabilizing charge surface without compensating metal ion or positive charge group. The single mutants where Ser residue is mutated to Arg in Hahellin-S80R and Hahellin-S41R show a marked decrease in negative surface charge potential as compared to Hahellin-wt and NMR-derived holo-Hahellin. In the double mutant Hahellin-S41R-S80R, there is substantial decrease in the negative surface charge potential as shown in Figure 6, suggesting that the mutation to positively charged Arg residue at both canonical motifs decreases the repulsive negative surface charge potential and facilitates its stabilization. Thus, the decrease in negative surface charge potential is the direct consequence of mutation with a positively charged Arg residue in place of Ser.

**Figure 6.**
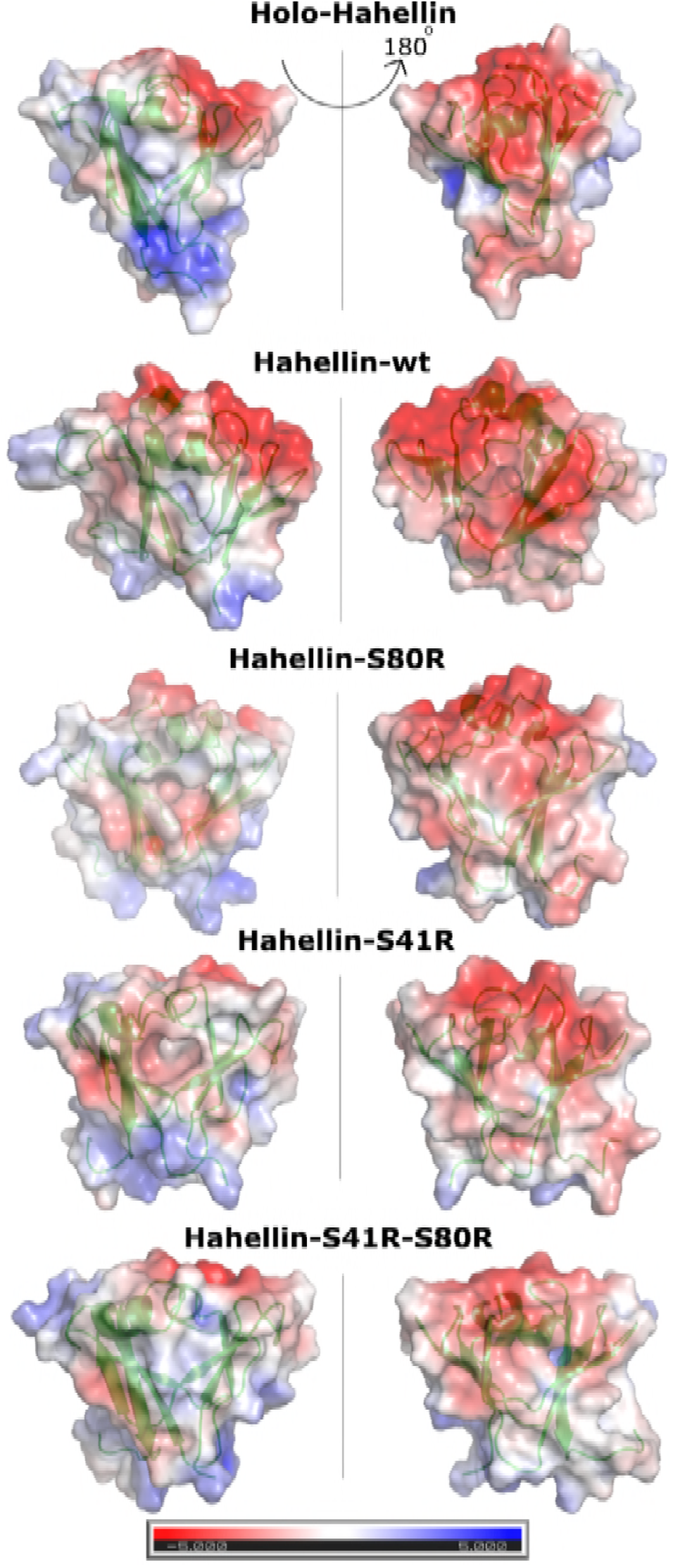
Surface charge potential on holo-Hahellin and the centroid structure of the most populated cluster of Hahellin-wt, Hahellin-S41R, Hahellin-S80R and Hahellin-S41R-S80R simulations. Red indicates negative charge potential, blue indicates positive charge potential and white colour indicate polar/non-polar charge potential. Centroid structures are shown in cartoon representation (green) and surface potential in transparency mode.

### New interactions formed at the Ca^2+^ binding canonical motifs of the mutants

Table 1 shows the hydrogen bonds and ionic interactions formed by the residue at 5^th^ position of the canonical Ca^2+^ binding motifs which are observed for more than 10% of the total simulation time and the same residues are also shown in Figure 7. Residue S41 in Hahellin-wt formed four hydrogen bonds (S41:sc-E9:mc, S41:mc-S42:sc, S41:mc-Y8:mc and S41:sc-N37:mc) while S80 shows only one hydrogen bond (S80:sc-D78:sc) (Table 1 and Figure 7). The mutated residue, R41 in Hahellin-S41R REMD simulation forms five interactions, out of which three are ionic interactions (R41:sc-D38:sc, R41:sc-D76:sc and R41:sc-D39-sc) and two are hydrogen bonding (R41:mc-Y8:mc and R41:sc-E9:mc). The R41:sc-D76:sc ionic interactions observed for 50.9% of the time, is a long-range interaction holding the β4 and β8 strands together. The S80 residue in Hahellin-S41R simulation did not form any persistent hydrogen bond which is observed for more than 10% of the time (Table 1). The S41 in Hahellin-S80R simulation shows only two hydrogen bonds (S41:sc-D39:sc and S41:sc-E9:mc) while the mutated residue R80 at the second Greek key forms two ionic interactions (R80:sc-D38:sc and R80:sc-D78:sc) and a single hydrogen bond (R80:mc-F54:mc). The interaction R80:sc-D38:sc is a long-range interaction between the loop (formed by the α-helix and β4-strand) and β8-strand, observed for 27.5% of the time which decreases the flexibility of the loop. Similar interaction is also observed in the crystal structure of a homologous protein, Clostrillin. In the double mutant Hahellin-S41R-S80R, six interactions (R41:sc-D38:sc, R41:sc-D39:sc, R41:sc-D76:sc, R41:mc-E9:mc, R41:sc-Y8:mc and R41:sc-E72:sc) are formed by R41 residue while two interactions (R80:mc-F54:mc and R80:sc-D38:sc) are formed by R80 residue (Table 1 and Figure 6). The ionic interaction, R41:sc-D76:sc is a long-range interaction observed for 57% of the time, connecting β4 and β8 strands together thus contributing to the overall βγ-crystallin stability while the rest two ionic interactions (R41:sc-D38:sc and R41:sc-D39:sc) are of short-range type, important for local turn stability. The double mutant forms maximum number of interactions as compared to rest of the simulations (Table 1 and Figure 4). Further, it can be noted that the mutant R41 in the first Greek key motif contributed significantly in terms of interactions compared to R80 in the second Greek key motif. Figure S2 to show special arrangement of the Ca^2+^ coordinating residues which are more compact in Hahellin-S41R-S80R similar to that of holo-Hahellin. So, we observed that several short-range and long-range interactions involving distinct β-strands and loops regions provide overall stability to the βγ-crystallin in the double mutant.

**Table 1.** Types and percentage of hydrogen bond/ionic interactions formed by the residue at 5^th^ position of both the canonical motifs. The hydrogen bond/ionic interactions having greater than 10% are shown in the table. Ionic interactions are indicated in bold.

| Hydrogen bond/ ionic interaction | Percentage (%) | Hydrogen bond/ ionic interaction | Percentage (%) |
| --- | --- | --- | --- |
| **Hahellin-wt** |  |  |  |
| **S41** |  | **S80** |  |
| S41:mc-Y8:mc | 12.3 | **S80:sc-D78:sc** | **15.3** |
| S41:sc-E9:mc | 25.2 |  |  |
| S41:mc-S42:sc | 11.0 |  |  |
| S41:sc-N37:mc | 10.6 |  |  |
| **Hahellin-S41R** |  |  |  |
| **S41R** |  | **S80** |  |
| R41:mc-Y8:mc | 14.1 |  |  |
| **R41:sc-D10:sc** | **24.5** |  |  |
| **R41:sc-D38:sc** | **48.9** |  |  |
| **R41:sc-D39:sc** | **22.9** |  |  |
| **R41:sc-D76:sc** | **50.9** |  |  |
| **Hahellin-S80R** |  |  |  |
| **S41** |  | **S80R** |  |
| S41:sc-E9:mc | 13.5 | R80:mc-F54:mc | 20.2 |
| **S41:sc-D39:sc** | **20.9** | **R80:sc-D38:sc\*** | **27.5** |
|  |  | **R80:sc-D78:sc** | **44.0** |
| **Hahellin-S41R-S80R** |  |  |  |
| **S41R** |  | **S80R** |  |
| R41:sc-Y8:mc | 19.9 | R80:mc-F54:mc | 26.6 |
| R41:mc-E9:mc | 19.3 | **R80:sc-D38:sc\*** | **31.3** |
| **R41:sc-D38:sc** | **43.8** |  |  |
| **R41:sc-D39:sc** | **45.5** |  |  |
| **R41:sc-E72:sc** | **14.5** |  |  |
| **R41:sc-D76:sc** | **57.0** |  |  |

\*Similar ionic interaction is observed in Clostrillin-T82R crystal structure where 5<sup>th</sup> residue of canonical Ca<sup>2+</sup> ligating Thr residue is mutated to Arg.

**Figure 7.**
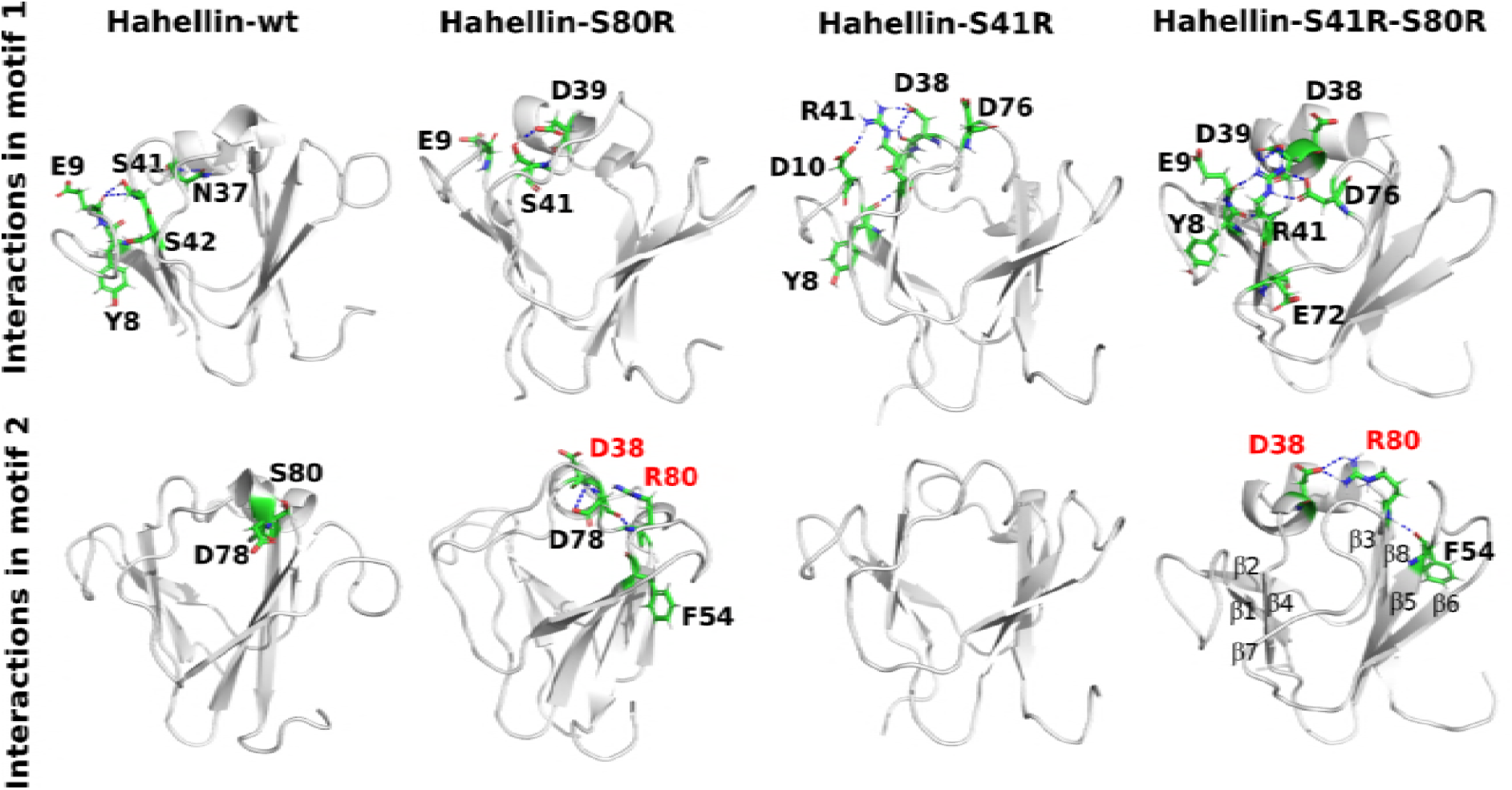
Residues at 5^th^ position of the canonical Ca^2+^ binding sites of both motif 1 and motif 2 and its interacting residues involved in hydrogen bonding and/or ionic interactions are shown in stick model. The labels in red indicates that they involved in ionic interaction (R80:sc-D38:sc) which was also observed in the crystal structure of a homologous βγ-crystallin, Clostrillin-S80R.

## Discussion

### Transformation of intrinsically disordered Hahellin into an ordered state is driven by several newly formed interactions

A typical Ca^2+^ binding βγ-crystallin domain consists of a double Greek key topology having two canonical Ca^2+^ binding motifs (N/D-N/D-X_1_-X_2_-S/T-S) where the first two positions have preference for Asn or Asp, the 3^rd^ and 4^th^ positions are occupied by any residue, while the 5^th^ position has preference for a Ser or Thr and the 6^th^ position has a Ser [22]. Mutation at the 5^th^ position of the canonical motif by Arg decreases and/or abolishes the Ca^2+^ binding affinity and makes it a disabled motif [22]. In the Ca^2+^ bound form, four (+x_mc_, -x_sc_, +y_mc_, +z_sc_) out of six or seven coordination sites are contributed by the polypeptide itself while the remaining sites are coordinated by water. There are two canonical Ca^2+^ binding motifs ^37^NDDLSS^42^ and ^76^DNDASS^81^ in Hahellin ligating two Ca^2+^ ions. Precisely, the Ca^2+^ ligating residues are E9(+x_mc_), N77(-x_sc_), D39 (+y_mc_) and S41 (+z_sc_) forming the site 1 and Q55(+x_mc_), D38(-x_sc_), D78(+y_mc_) and S80 (+z_sc_) forming the site 2 based on sequence comparison with other known βγ-crystallins[16]. The residues at +x_mc_ and +y_mc_ coordinates through main-chain carbonyls while residues at -x_sc_ and +z_sc_ coordinate through sidechains. The Ca^2+^ binding pocket is further stabilized by hydrogen bonding between the first residue of the motif 1 (^37^NDDLSS^42^) and the fifth residue of motif 2 (^76^DNDASS^81^) [16]. βγ-crystallin superfamily includes proteins of diverse domain stability. Many members have high intrinsic domain stability while others require Ca^2+^ to gain further stability [17]. There are certain highly disordered proteins such as Hahellin, Caulollin [18] and Yersinia crystallin [19] that undergo a large conformational change to βγ-crystallin upon binding to Ca^2+^. In the present study, we observed that Hahellin samples intrinsically disordered states in the absence of Ca^2+^. However, upon mutation at 5^th^ position of the canonical motif from a Ser to Arg in Hahellin-S41R and Hahellin-S80R, there is significant decrease in the disordered population and concomitant increase in the ordered conformations. On the other hand, remarkable transformation was observed in the case of double mutant where, Hahellin-S41R-S80R samples mostly ordered conformations like that of Ca^2+^ bound Hahellin. Upon detailed examination, we find that there are several new long-range ionic and hydrogen bonding interactions formed by the mutant residue at the Ca^2+^ binding sites which decrease the loops flexibility and increase the overall stability of the βγ-crystallin fold (Table 1 and Figure 7). The residues Y8 and E9 belong to β1-strand, residues D38 and D39 located in the loop close to β4-strand, while residue D76 is located in the inter β-strand loop formed by β7 and β8 strands. The mutated residue, R41 interacts with Y8, E9, D38, D39 and D76 residues intermittently and acts as a junction point to hold the β-strands β1, β4, β7 and the connecting loops in place. Since the β-strands β1 and β2 form a β-hairpin, the first Greek key motif formed by β1, β2, β4, β7 strands and loops becomes more rigid. On the other hand, stability of the second Greek key motif is achieved by the formation additional new ionic and hydrogen bonding interactions involving the mutated R80 residue in Hahellin-S80R and in double mutant Hahellin-S41R-S80R (Table 1 and Figure 7). The residue R80 is close to β8-strand of second Greek key, D38 residue is in the loop connecting the flexible α-helix and the β4-strand of first Greek key and the residue F54 is in β5-strand of second Greek key. The strands β8 and β5 in the second Greek key have several inter-β-stand interactions in addition to R80:mc-F54:mc hydrogen bond and long-range ionic interactions (R80:sc-D38:sc) which stabilize the second motif (Figure 7). Besides, there are bridging residues connecting the first motif and the second motif. D38 is one such residue which forms ionic interactions with mutant R41 belonging to the first motif as well as with R80 belonging to the second motif in the double mutant. D76 is another such bridging residue which does not interact with the mutant residue but forms hydrogen bonding interaction with N77 of motif 2 Ca^2+^ ligating site and ionic interaction with R44, located close to the motif 1. The bridging residues play an important role in interlocking the two motifs close together to make it more compact like that of holo-Hahellin. Thus, several newly formed interactions within and between the motifs along with the existing ones provides overall stability to the βγ-crystallin domain without Ca^2+^, as observed in our simulations.

### Overall stabilization of βγ-crystallin domain in Hahellin is largely dictated by rigidification of first canonical motif

We observed in our simulations that Hahellin-S41R and Hahellin-S41R-S80R sample more homogenous ensembles whereas Hahellin-S80R samples partially unfolded conformations and Hahellin-wt samples largely disordered conformations in the absence of Ca^2+^. In the disordered conformations of Hahellin-wt and Hahellin-S80R, the N-terminal Greek key is found to be more disordered while the C-terminal Greek key is partially ordered which is evident from the Cα RMSF plot and native contact analysis (Figure 4b and 6). Mutation of Ser to Arg at the first canonical motif resulted in more ordered conformations due to formation of greater number of ionic and hydrogen bonding interactions (Table 1 and Figure 7). The C-terminal Greek key is inherently ordered and the mutation (S80R) at the second canonical motif does not show drastic difference in terms of newly formed interactions. Thus, overall stabilization of βγ-crystallin domain in Hahellin is largely dictated by rigidification of the first canonical motif which is achieved by S41R mutation.

### Increase in domain stability due to mutation of Ser to Arg at 5^th^ position of the canonical N/D-N/D-X_1_-X_2_-S/T-S motif in Hahellin is consistent with other experimental studies

Although the βγ-crystallins have a characteristic double Greek key motif their Ca^2+^ binding ability varies in the different organisms. Most of the microbial βγ-crystallins possess canonical Ca^2+^ binding motifs, whereas, the vertebrate eye-lens crystallins possess non-canonical motifs. The occurrence of non-canonical motif results in decrease and/or abolition of Ca^2+^ binding affinity. Domain stability in microbial βγ-crystallins increased further upon binding to Ca^2+^. Although, the non-canonical βγ-crystallins motifs have decreased affinity for Ca^2+^, they possess high intrinsic domain stability which is established from a comparative study on four diverse βγ-crystalllin [17]. A selective mutation of Ser/Thr to Arg at 5^th^ position of the canonical motif was carried out on Vibrillin (motif1: NDTLSS and motif2: NDVISS), Flavollin (motif1: NDDISS and motif2: NDKVSS), Clostrillin (motif1: NDWMTS and motif2: NDKMTS) and Centrillin (motif1: NDRARS and motif2: FRQISS) and their stabilities were accessed by GdmCl-induced unfolding [17]. Centrillin has one non-canonical (NDRARS) and one canonical (FRQISS) motifs whereas other three possess only canonical motifs. By having one non-canonical motif where 5^th^ position has a natural Arg, the stability of Centrillin was observed to be greater than the rest three βγ-crystallins.

In another study, mutation was done individually and together at 5^th^ position of both the canonical Ca^2+^ binding motifs of Clostrillin from Thr to Arg. These mutations increase the domain stability, make it comparable to that of holo-Clostrillin with concomitant decrease in Ca^2+^ binding affinity [22]. In the crystal structure of Clostrillin-T82R which could only be crystallized, R82 which is the 5^th^ residue (+z_sc_) of second canonical motif forms hydrogen bonds and ionic interactions with D38 which is the 2^nd^ residue of first canonical motif [22]. Similar ionic interactions (R80:sc-D38:sc) between D38 (2^nd^ residue of first canonical motif) and S80 (5^th^ residue of second canonical motif) are observed in both Hahellin-S80R and Hahellin-S41R-S80R simulations. In the absence of Ca^2+^, this along with other hydrogen bonds and ionic interactions as mentioned in the Results stabilize the βγ-crystallin fold. Arg having a branched sidechain facilitates the intra-motif and inter-motif interactions which is the sole cause of overall stabilization of the βγ-crystallin domain.

### Plausible cause for the decrease in Ca^2+^ binding affinity upon mutation

There are several studies which establish that the Ca^2+^ binding affinity is decreased upon mutation to Arg at 5^th^ position of the canonical Ca^2+^ binding site [17, 22]. One reason could be the +ve charge on Arg sidechain which creates a repulsive potential for the incoming Ca^2+^ ions with similar charge. The second reason could be due to involvement of Ca^2+^ ligating residues in several interactions as shown in Table 1 which distort the geometry required for Ca^2+^ binding. The branched sidechain of R41 in motif 1 forms interactions with E9 carbonyl oxygen and D39 sidechain which provides ligating sites to the Ca^2+^ in holo-form. Similarly, R80 in the motif 2 distorts the Ca^2+^ coordinating geometry due to R80:sc-D38:sc interaction. Because of these newly formed interaction by D38(-x_sc_), the second Ca^2+^ ligating site is lost. Therefore, upon mutation ligation sites are unavailable for coordination with Ca^2+^ which add-in in decreasing the affinity for Ca^2+^. Similar, decrease in Ca^2+^ binding affinity is also reported for Clostrillin-T41R-T82R [22].

## Conclusion

We observed disordered states as well as ordered state of Hahellin which are modulated by mutation at 5^th^ position of the canonical motifs. Thus, the choice of residue at the canonical motifs has remarkable impact on the structural properties of Hahellin. In particular the Arg residue at 5^th^ position of the canonical motif proved to be consistently important for the structural integrity of βγ–crystallin domain. Arg at 5^th^ position not only neutralizes the negative surface charge potential but also forms several long-range and short-range ionic and hydrogen bonding interactions through its branched sidechain with adjacent β–strands and loop regions altering the required Ca^2+^ binding geometry by preoccupying the Ca^2+^ binding sites through formation several interactions. The newly formed interactions form by the mutant residue causes rigidification within and between the two motifs to make it more compact like that of Ca^2+^ bound βγ–crystallin. This is the reason why it shows low affinity for Ca^2+^ with concomitant high stability of the βγ–crystallin domain. Mutation at first canonical Ca^2+^ binding motif (S41R) observe to stabilize the βγ–crystallin domain better than the mutation (S80R) alone at the second Greek key motif. The study thus helps in identifying the crucial interactions required for transition from disordered-to-order state and vice-versa.

## Methods

### Starting structures

The starting structures used for the simulations were derived from NMR which is Ca^2+^ bound form of Hahellin having a well ordered βγ-crystallin fold [20]. For the wild type simulation, NMR derived 3D structure without Ca^2+^ was used and is abbreviated as Hahellin-wt. The mutants of Hahellin were generated by mutating S41 and S80 to Arg which occupy 5^th^ position of the first and second canonical Ca^2+^ binding motifs (N/D-N/D-X_1_-X_2_-S/T-S) respectively as shown in the Figure S3 [22]. These mutants thus generated are abbreviated as Hahellin-S41R, Hahellin-S80R and Hahellin-S41R-S80R (Figure S3).

### Replica Exchange Molecular Dynamics (REMD) simulation

REMD simulations were performed on the wild type protein (Hahellin-wt) and on the three mutants (Hahellin-S41R, Hahellin-S80R and Hahellin-S41R-S80R) by keeping all the parameters same. REMD is an efficient method for enhanced conformational sampling. In this method, instead of a single molecular dynamics run, multiple parallel runs were initiated at select replica temperatures. The temperatures were selected from an exponential distribution so that the interval between the temperatures is relatively more at higher temperature to keep the exchange rate constant across all the replicas. The conformations (coordinates) were exchanged at regular intervals between the neighboring replica temperatures following the Metropolis algorithm proposed by Sugita and Okamoto [27]. Periodic evaluation (every 2 ps) of Metropolis algorithm was performed in order to exchange coordinates between the neighbouring replicas. When the condition was satisfied, the coordinates of the neighbouring replicas were swapped, and the exchange attempt was considered successful. The velocity of the target replica was then rescaled to the target temperature. In this study, we used sixteen replicas temperatures such as 281.85, 285.39, 288.98, 292.60, 296.28, 300.00, 303.77, 307.58, 311.44, 315.36, 319.32, 323.33, 327.39, 331.50, 335.66 and 339.88 K with an exchange probability of 60%.

### Simulation details

AMBER 12 molecular modeling package was used to perform all REMD simulations [28]. The force-field used was AMBER FF03 [29, 30]. Implicit solvent with generalized Born solvation model was used to treat the solvent [31] under NVT ensembles [32]. The topology, coordinates and force-field parameters were generated using the *tleap* program [28]. The side-chains of the polar residues were adjusted to confirm neutral pH conditions. Initially, the structure was subjected to energy minimization for 2000 cycles and the first 1000 cycles were performed by steepest decent energy minimization and the next 1000 cycles were subjected to conjugate gradient minimization to remove van-der-Waals contacts of high potential energy. The chirality constraints were generated on the minimized structure to prevent unwanted rotation around the peptide bonds, which might occur at higher replica temperatures during the simulation. SHAKE algorithm was used to constrain bond stretching freedom of all bonds involving hydrogen atoms [33]. The non-bonded van-der-Waals cut-off was set at 16 Å. The temperature of each replica was maintained by weak coupling to the Langevin thermostat with a collision frequency of 1 ps^−1^. The system was equilibrated for 200 ps during which the temperature of each replica was gradually increased from 0 to the target temperature. Following the equilibration, REMD simulation was initiated for each system with an integration step of 2 fs, during which the exchange attempts were made every 2 ps between the neighboring replica temperatures. Parallel REMD run was performed using *Multisander* [28] program which uses the *Verlet* integration algorithm. The resultant coordinate and output files from all replicas were saved every 2 ps. Each REMD simulation was performed for 100 ns per replica. The program *cpptraj* was used to process and filter the trajectories [28] which passes through different replica temperatures due to exchange. For the data analysis, the trajectory corresponding to 300 K temperature was filtered for each simulation. REMD simulations were run in the high performance computing facility of TIFR Centre for Interdisciplinary Sciences.

### Convergence

The REMD trajectory of individual simulation was divided into 10 ns overlapping segments and the numbers of clusters as a function of time segment were plotted. 5 Å C^α^ RMSD cutoff with respect to the starting structure was used for the cluster analysis of each segment (Figure S4). The plot showed a gradual increase in the number of clusters till it reaches the time segment of 0-60 ns. After which the number of clusters were seemingly constant till the last time segment. Therefore, we considered 60 to 100 ns as equilibration period and performed all the analyses corresponding to this time segment.

### Analyses

The analyses of REMD simulations were performed by monitoring the following conformational parameters. These are (i) radius of gyration (Rg) of the backbone atoms, (ii) the root-mean-square deviation (RMSD) of C^α^ atoms with respect to the NMR derived 3D structure of holo-Hahellin, (iii) free energy landscape with Rg and RMSD as the coordinates, (iv) clustering and network layout (details given below), (v) the percentage of native contact was determined for the centroid structures by evaluating the number of non-local contacts with respect to holo-Hahellin within a cut-off radius of 6.5 Å, (vi) root-mean-square-fluctuation of Cα atoms, (vii) surface charge distribution on the centroid structure of the most populated cluster, (viii) ionic interaction was determined by finding distance between the positive and negative charged sidechain atoms which should be less than 4 Å (ix) hydrogen bond was determined as defined by Visual Molecular Dynamics (VMD) [34]. VMD software was used for visualization of each trajectory and to perform structural analyses by using *tcl* scripting implemented therein. Matlab (http://www.mathworks.com) and Xmgrace (http://plasma-gate.weizmann.ac.il/Grace) were also used in the structural analyses and for plotting the graphs. The Amber tool *cpptraj* was also used in some cases [35].

### Free energy landscape

The free energy landscape was built using backbone Rg and C^α^ RMSD for the equilibrium time segment of 60 to 100 ns. The Rg-RMSD plane was divided into 20 × 20 grid cells, resulting into 400 cells. The free energies were then determined for each cell with respect to reference cell containing maximum number of points by evaluating ∆A_ref→i_ = -RT ln(N_i_/N_ref_), where N_i_ and N_ref_ are the number of points in i^th^ and reference cell, respectively, R is the ideal gas constant and T is the temperature [36, 37]. We filtered conformations corresponding to the free energy minima and clustered these conformations to find the centroid structure, which explained the conformational features of the peptide on the selected region of the free energy landscape.

### Clustering and network layout

Clustering and network layout method was used to segregate structures into distinct clusters. In this method, a pairwise C^α^ RMSD matrix was built and a pairwise RMSD cutoff was chosen to form the network layout. A network layout was made up of nodes and links. Each structure in the network layout represented a node. The links were established based on the chosen pairwise C^α^ RMSD cutoff, which was determined following Ahlstrom et al.[38]. For this, we employed profuse-force-directed layout [39] which comes with Cytoscape software [40]. For clustering the structures and to generate centroid structure corresponding to each cluster, an average linkage algorithm[41] in *Matlab* was used. The centroid structures thus generated represent the central conformation of a given cluster.

## Acknowledgments

S.P. acknowledges support from DST for WOS-A with grant number (No. SR/WOS-A/CS-143/2017). We thank Yogendra Sharma (CCMB) for critically going through the manuscript.

